# *In Silico* Design of APOE ɛ4 Interaction Inhibitor Peptides for Alzheimer’s Disease

**DOI:** 10.1101/2025.09.03.673973

**Authors:** Jaeho ji, Eunjeong Han, Jongham Park, Ahrum Son, Hyunsoo Kim

## Abstract

**Background:** Protein-protein interactions (PPIs) are essential for cellular functions, and their aberrant formation contributes to neurodegenerative diseases. Alzheimer’s disease (AD), characterized by its complex pathogenesis, poses an increasing societal burden with population aging. The APOE ɛ4 allele represents the strongest genetic risk factor for late-onset AD, yet its pathological mechanisms remain incompletely understood.

**Methods:** We employed artificial intelligence-driven peptide design to elucidate the pathological interaction between APOE ɛ4 and amyloid precursor protein (APP). Using advanced AI algorithms, we identified critical binding interfaces and designed mimetic peptides targeting the APOE ɛ4–APP interaction site. Peptide efficacy was evaluated through comprehensive molecular dynamics simulations.

**Results:** Our analysis revealed key residues mediating APOE ɛ4–APP binding. The designed inhibitory peptides demonstrated stable interaction with target sites, favorable binding energetics, and sustained structural integrity throughout simulations. Lead candidate effectively disrupted APOE ɛ4– APP complex formation *in silico*.

**Conclusion:** This study presents a novel AI-powered approach for developing PPI-targeted therapeutics against AD. Our computationally validated peptide inhibitors offer promising therapeutic candidates that warrant experimental validation. These findings demonstrate the potential of integrating artificial intelligence with structural biology for accelerating drug discovery in neurodegenerative diseases.

## Introduction

Protein-protein interactions (PPIs) constitute the molecular foundation of cellular life, orchestrating essential biological processes ranging from signal transduction and metabolic regulation to cell survival and programmed death [1–6]. While properly regulated PPIs ensure normal protein folding and functional execution, their dysregulation has emerged as a critical factor in numerous pathological conditions, most notably in neurodegenerative disorders such as Alzheimer’s disease [7–9].

Alzheimer’s disease (AD) stands as the predominant from of dementia worldwide, presenting a formidable challenge to modern medicine through its progressive cognitive decline, memory deterioration, and behavioral alterations [1–3, 5, 6, 10]. Despite decades of intensive research, the complex pathogenesis of AD continues to elude complete understanding. The diseases manifests through distinctive pathological signatures: extracellular neuritic plaques composed of amyloid-β (Aβ) peptide fibrils, intracellular neurofibrillary tangles of hyperphosphorylated tau protein, and marked deficits in neurotransmitter systems [11, 12].

A pivotal breakthrough in AD research came in 1993 with the identification of apolipoprotein E (APOE) as a major genetic risk factor [13–16]. This 317-amino acid glycoprotein, functioning primarily as cholesterol transporter in the brain, exists in three allelic variants–ɛ2, ɛ3, and ɛ4– distinguished by cysteine and arginine substitutions at positions 130 and 176 (ɛ2: Cys130/Cys176; ɛ3: Cys130/Arg176; ɛ4: Arg130/Arg176) [17–21]. Though unable to cross the blood-brain barrier, APOE is abundantly produced within the central nervous system by astrocytes, activated microglia, vascular wall cells, choroid plexus cells, and stressed neurons. These seemingly minor amino acid variations profoundly impact APOE’s structural conformation, subsequently affecting its lipid-binding capacity, receptor interactions, oligomerization patterns, and overall stability [21–26].

The functional consequences of APOE isoform variations extend far beyond simple structural differences. Research has revealed that APOE ɛ4 preferentially associates with triglyceride-rich particles, such as very low-density lipoprotein (VLDL), whereas ɛ2 predominantly binds to high-density lipoprotein (HDL) particles [21, 27, 28]. This differential lipoprotein association has profound implications for Aβ metabolism. Lipidated APOE promotes Aβ degradation through the ATP-binding cassette transporter A1 (ABCA1) pathway; however, APOE ɛ4’s reduced lipid transport efficacy compared to ɛ2 significantly impairs this clearance mechanism [21, 29, 30]. Notably, peptide mimetics that upregulate ABCA1 have shown promise in enhancing APOE ɛ4 lipidation, thereby promoting Aβ clearance and ameliorating tau-related cognitive deficits in preclinical models [21, 31].

The pathological impact of APOE ɛ4 extends to microglial function, a critical component of the brain’s immune response. Studies demonstrate that APOE ɛ4 is markedly less effective than the canonical ɛ3 variant in activating microglial responses to Aβ accumulation [21, 32]. This impairment appears to be mediated through the triggering receptor expressed on myeloid cells 2 (TREM2), a microglial-specific receptor whose interaction with APOE depends on both isoform type and lipidation status [21, 33–35]. The reduced binding affinity between APOE ɛ4 and TREM2 may thus contribute to compromised microglial surveillance and clearance functions, further exacerbating AD pathology [21].

The differential risk profiles of APOE variants are striking: ε4 increases AD risk 3-fold in heterozygotes and 15-fold in homozygotes, while ε2 confers protection, reducing disease risk by approximately 50% and delaying Aβ deposition. APOE ε4’s pathogenic mechanisms encompass impaired Aβ clearance, reduced membrane lipid recycling efficiency, compromised neuronal repair capacity, and enhanced Aβ fibrillation and plaque formation [13, 16, 21, 36–49]. These observations position APOE-targeted interventions, particularly those modulating APOE-mediated inflammatory responses, as promising therapeutic strategies.

Recent technological advances have opened new avenues for therapeutic development. The emergence of generative artificial intelligence algorithms such as ProteinMPNN and RFdiffusion has revolutionized de novo protein and peptide design, enabling the creation of novel therapeutic molecules with unprecedented precision and specificity [50, 51]. These generative models can design entirely new protein sequences and structures tailored to bind specific targets, moving beyond mere structure prediction to active molecular engineering. By leveraging these tools to identify and target critical amino acids at protein interfaces—termed “hot spots“—researchers can now design bespoke peptide inhibitors with optimized binding properties.

In this study, we leverage these generative AI approaches to design and optimize peptide inhibitors targeting the pathological interactions between APOE ε4 and its partner proteins. Our objective is to elucidate the molecular basis of AD-specific aberrant protein-protein interactions and employ generative design algorithms to create targeted peptide therapeutics that disrupt these pathological interactions. By focusing on interface hotspots and utilizing advanced generative models, we aim to develop APOE ε4-specific therapeutic peptides with enhanced binding affinity and specificity that could potentially modify disease progression. This approach represents a novel paradigm in AD therapeutic development, offering the potential for precision-designed interventions tailored to individual genetic risk profiles.

## Materials and Methods

### Protein Interactome Structure Prediction

Structural models of protein complexes involving APOE allele-specific interacting partners were generated using AlphaFold3 [52]. These predictions provided the foundation for subsequent structural analyses and inhibitor design.

### Structural Analysis of Protein Interactomes

Comprehensive structural analysis and interaction of protein-protein interaction interfaces were performed using a combination of computational approaches. Initial visualization and interaction site mapping were conducted using PyMOL (Schrödinger, LLC). We also conducted solvent-accessible surface area (SASA) calculations using GROMACS [53]. SASA analysis quantifies the surface area of biomolecules accessible to solvent molecules, providing insights into conformational changes upon protein-protein interactions [54–56].

We performed comparative SASA analysis between isolated APOE allele structures and their corresponding protein complexes. Regions exhibiting significant changes in surface area upon complex formation were identified as putative interaction sites. By integrating results from both PyMOL visualization and SASA calculations, we identified key interaction hotspots and selected proteins displaying allele-specific differences in interaction pattern as primary targets for inhibitor development.

### Peptide Inhibitor Design

Peptide inhibitors targeting identified interaction hotspots were designed through a multi-step computational pipeline. Initial backbone scaffolds were generated using RFdiffusion to specifically target the characterized interaction sites within the protein interactome [50]. Subsequently, functional group optimization was performed using ProteinMPNN coupled with FASTRELAX protocols to develop peptides capable of disrupting protein-protein interactions [51, 57].

### Blood-Brain Barrier Permeability Prediction

Effective therapeutic intervention in Alzheimer’s disease requires that designed peptide inhibitors cross the blood-brain barrier (BBB) to reach their targets within the central nervous system. We evaluated BBB permeability of our peptide libraries using the DeepB3P3 algorithm, a deep learning-based predictor trained on experimental permeability data [58].

### Molecular Docking Simulations

To assess the binding potential of BBB-permeable peptides, we performed molecular docking simulations using HADDOCK 2.4 [59, 60]. These simulations provided insights into the potential therapeutic efficacy of peptides as Alzheimer’s disease interventions.

### Binding Affinity Prediction

Theoretical binding affinities were calculated for both native APOE allele-protein complexes and designed peptide-target protein interactions using PRODIGY [4]. These predictions enabled comparative assessment of inhibitor potency across different APOE alleles.

### Stability and Dynamic Analysis

Structural stability, flexibility, and dynamic properties of designed peptides were comprehensively evaluated using multiple computational metrics. Radius of gyration (Rg) was calculated to assess overall structural compactness and stability [61]. Root mean square fluctuation (RMSF) analysis quantified residue-level flexibility and local stability [62]. Dynamic cross-correlation matrix (DCCM) analysis was employed to identify correlated motions between residues, providing insights into functional dynamics and interaction mechanisms [63]. All calculations were performed using the MDAnalysis package in Python [64, 65].

## Results

### APOE Allele Structure Comparison

The computational workflow employed in this study enabled systematic prediction and characterization of APOE allele structures and their interaction networks in the context of Alzheimer’s disease pathogenesis (**Figure 1**). This integrated approach facilitated the identification of allele-specific interaction interfaces that subsequently served as templates for designing peptide-based inhibitors.

**Figure 1.**
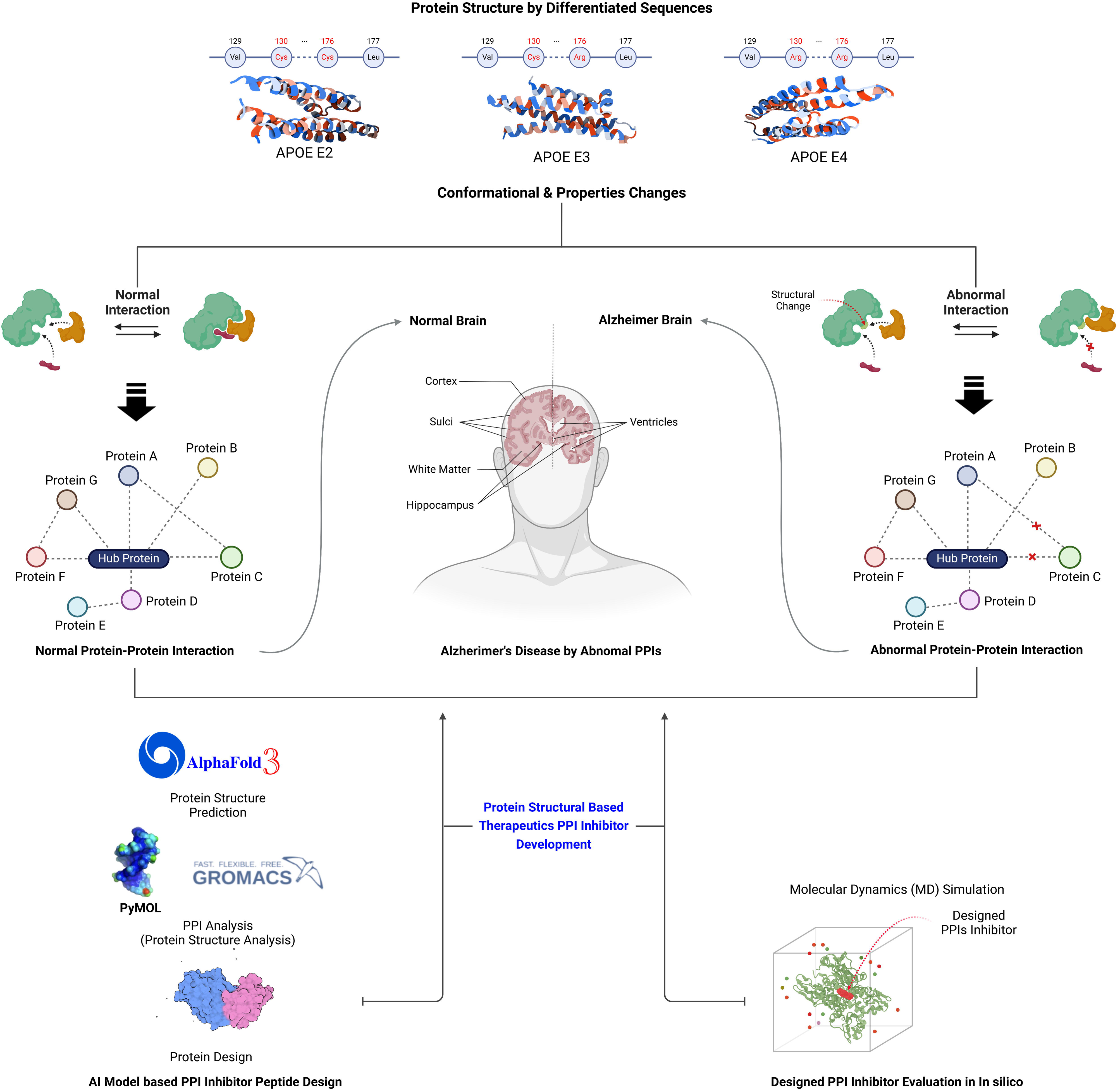
Computational workflow for structure-based peptide inhibitor design targeting APOE allele-specific interactions in Alzheimer’s disease. Schematic representation of the integrated computational pipeline employed for designing allele-selective peptide inhibitors. The workflow begins with the amino acid sequences of APOE isoforms (ε2, ε3, and ε4), which differ at positions 112 and 158. These sequence variations lead to differential protein-protein interactions associated with Alzheimer’s disease pathogenesis. Using AlphaFold3 and complementary structural analysis tools (PyMOL and SASA), we predicted the three-dimensional structures of APOE allele complexes with disease-relevant interacting partners (LDLR, VLDLR, ABCA1, TREM2, and APP) and identified allele-specific interaction interfaces. The characterized interaction hotspots served as target sites for rational peptide design using RFdiffusion and ProteinMPNN. Designed peptides were subsequently evaluated through multiple computational filters including BBB permeability prediction (DeepB3P3), molecular docking (HADDOCK), and stability assessments (RMSF, Rg, DCCM) to identify lead candidates capable of selectively disrupting pathological APOE ε4-mediated interactions.

To investigate whether the amino acid substitutions distinguishing APOE ε2 (Cys112/Cys158), ε3 (Cys112/Arg158), and ε4 (Arg112/Arg158) variants induce conformational changes affecting protein function, we first analyzed the predicted structures of individual APOE alleles. The AlphaFold3 predictions yielded pTM (predicted Template Modeling) scores of 0.51, 0.52, and 0.51 for APOE ε2, ε3, and ε4, respectively. While these moderate confidence scores reflect the limited availability of experimentally determined full-length APOE structures in existing databases, they indicate that the predicted overall fold topology is likely reliable. (**Supplemental Fig.1**) Notably, the regions encompassing the allele-defining residues at positions 112 and 158 demonstrated consistently high prediction confidence across all three variants, supporting the validity of subsequent comparative analyses.

Structural superposition of the three APOE allele models revealed remarkable conservation of the overall protein architecture (**Figure 2**). The N-terminal receptor-binding domain (residues 1-191) and C-terminal lipid-binding domain (residues 201-299) maintained their characteristic four-helix bundle conformations across all variants. Minor variations were observed primarily in loop regions and side-chain orientations, particularly in the vicinity of the polymorphic sites. The root mean square deviation (RMSD) values between aligned backbone atoms ranged from 0.8-1.2 Å, confirming the absence of major conformational changes attributable to the amino acid substitutions.

**Figure 2.**
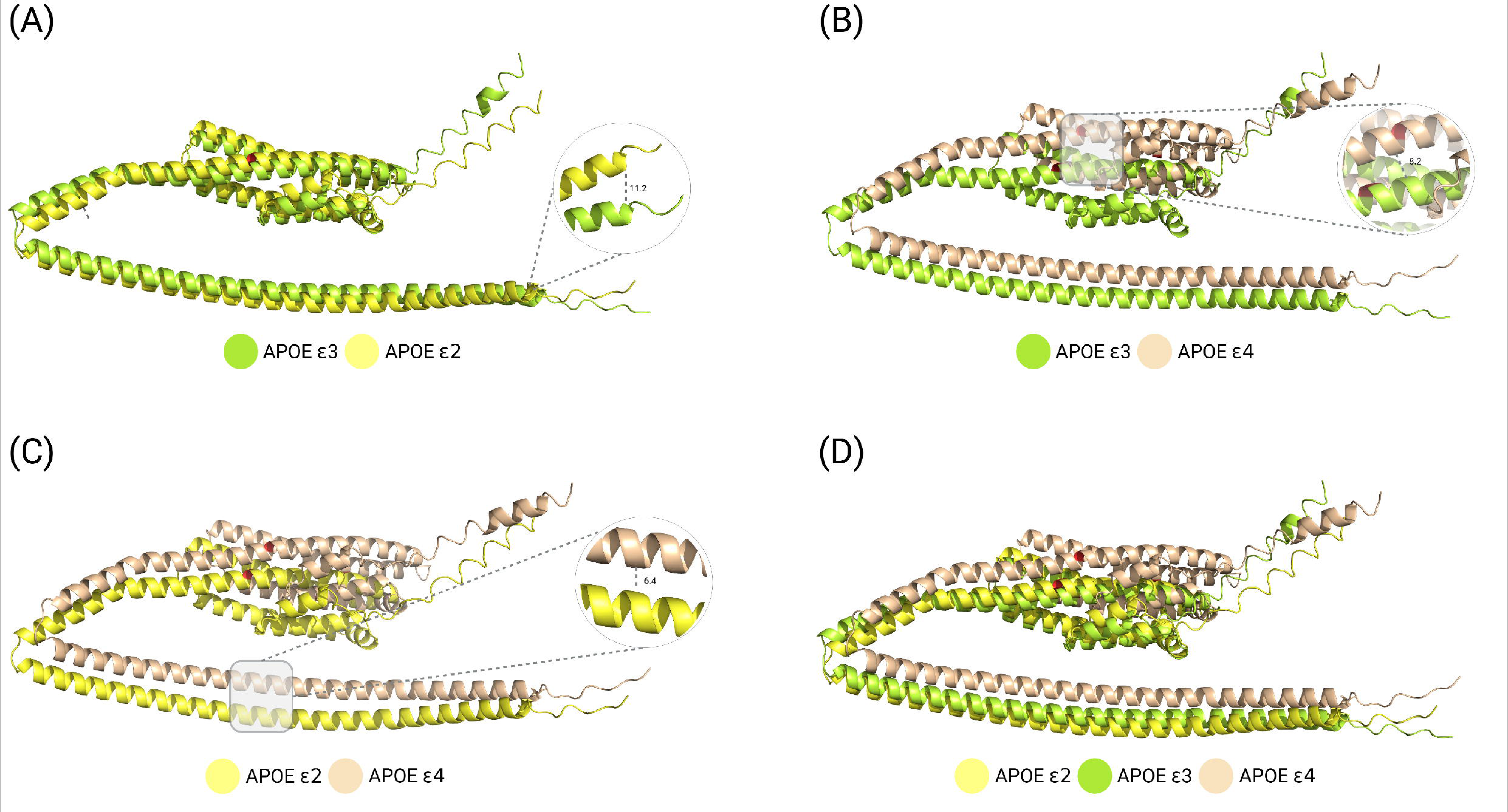
Structural comparison of APOE isoform predictions reveals conserved architecture across allelic variants. (A-C) AlphaFold3-generated structural models of APOE isoforms with corresponding confidence metrics. (A) APOE ε2 (Cys112/Cys158) structure colored by pLDDT confidence scores. (B) APOE ε3 (Cys112/Arg158) structure with confidence mapping. (C) APOE ε4 (Arg112/Arg158) structure displaying predicted confidence levels. Color gradients represent prediction confidence: blue (very high, pLDDT >90), cyan (high, pLDDT 70-90), yellow (moderate, pLDDT 50-70), and orange (low, pLDDT <50). (D) Structural superposition of all three APOE isoforms following sequence-based alignment. The overlay demonstrates highly conserved tertiary structure across variants, with RMSD values between aligned backbone atoms ranging from 0.8-1.2 Å. Despite amino acid substitutions at positions 112 and 158, no significant conformational changes are observed in the isolated protein structures, suggesting that functional differences between isoforms may primarily manifest through altered protein-protein interactions rather than intrinsic structural variations.

Despite the structural similarity observed in isolated APOE proteins, we hypothesized that allele-specific differences might manifest more prominently in the context of protein-protein interactions. Therefore, we proceeded to predict and analyze the complex structures of APOE variants with their respective interacting partners to elucidate how these subtle structural variations influence binding interfaces and interaction dynamics. This comparative analysis of protein complexes aimed to reveal the molecular basis underlying the differential pathological associations of APOE alleles in Alzheimer’s disease.

### APOE Protein Interactions in Alzheimer’s Disease Pathogenesis

The APOE protein engages in multiple interactions with key proteins implicated in Alzheimer’s disease pathology, including LDLR, VLDLR, ABCA1, TREM2, and APP. Notably, APOE isoforms exhibit distinct lipid particle associations, with APOE ε2 predominantly found in HDL-rich particles, whereas APOE ε4 primarily associates with VLDL-rich particles, facilitating differential interactions with VLDLR and LDLR proteins [20, 66]. The ABCA1 protein, a critical regulator of APOE lipidation, shows reduced membrane recycling in the presence of APOE ε4, resulting in increased ABCA1 aggregation and consequently decreased APOE ε4 lipidation [67]. These alterations compromise β-amyloid (Aβ) degradation capacity and contribute to Alzheimer’s disease pathogenesis.

TREM2, another crucial player in disease pathology, exhibits lipidation-dependent binding affinity for APOE proteins, with APOE ε4 showing reduced interaction that impairs microglial function [21, 34, 35]. Additionally, the amyloid precursor protein (APP), when aberrantly cleaved, generates Aβ peptides that accumulate to form characteristic amyloid plaques. APOE ε4 accelerates this pathological cascade by disrupting Aβ metabolism and clearance mechanisms [68, 69].

### Structural Prediction of APOE-Protein Complexes

To elucidate the structural basis underlying allele-specific differences in protein interactions, we employed AlphaFold3 to predict the three-dimensional structures of APOE variant complexes with their interacting partners. The prediction confidence scores and subsequent structural analyses revealed distinct patterns of interaction across different APOE alleles.

The structural predictions for APOE-LDLR complexes yielded pTM scores of 0.43, 0.45, and 0.48 for ε2, ε3, and ε4 variants, respectively (**Supplemental Fig.2**). Structural alignment based on LDLR revealed distinct binding poses across the three alleles. Using PyMOL visualization, we identified potential interaction residues with inter-residue distances below 5 Å (**Supplemental Table 1**). SASA analysis revealed significant surface area changes upon complex formation. The APOE ε2-LDLR complex showed differences in 31 residues (mean: 0.399, variance: 0.104, SD: 0.323), while the ε3-LDLR complex exhibited changes in 45 residues (mean: 0.384, variance: 0.082, SD: 0.287). The ε4-LDLR complex demonstrated alterations in 41 residues (mean: 0.355, variance: 0.070, SD: 0.264) (**Supplemental Fig.3A & Supplemental Table 1**). Distance measurements between residues with altered surface areas identified 11, 16, and 15 interacting residues for the ε2, ε3, and ε4 complexes, respectively.

**Figure 3.**
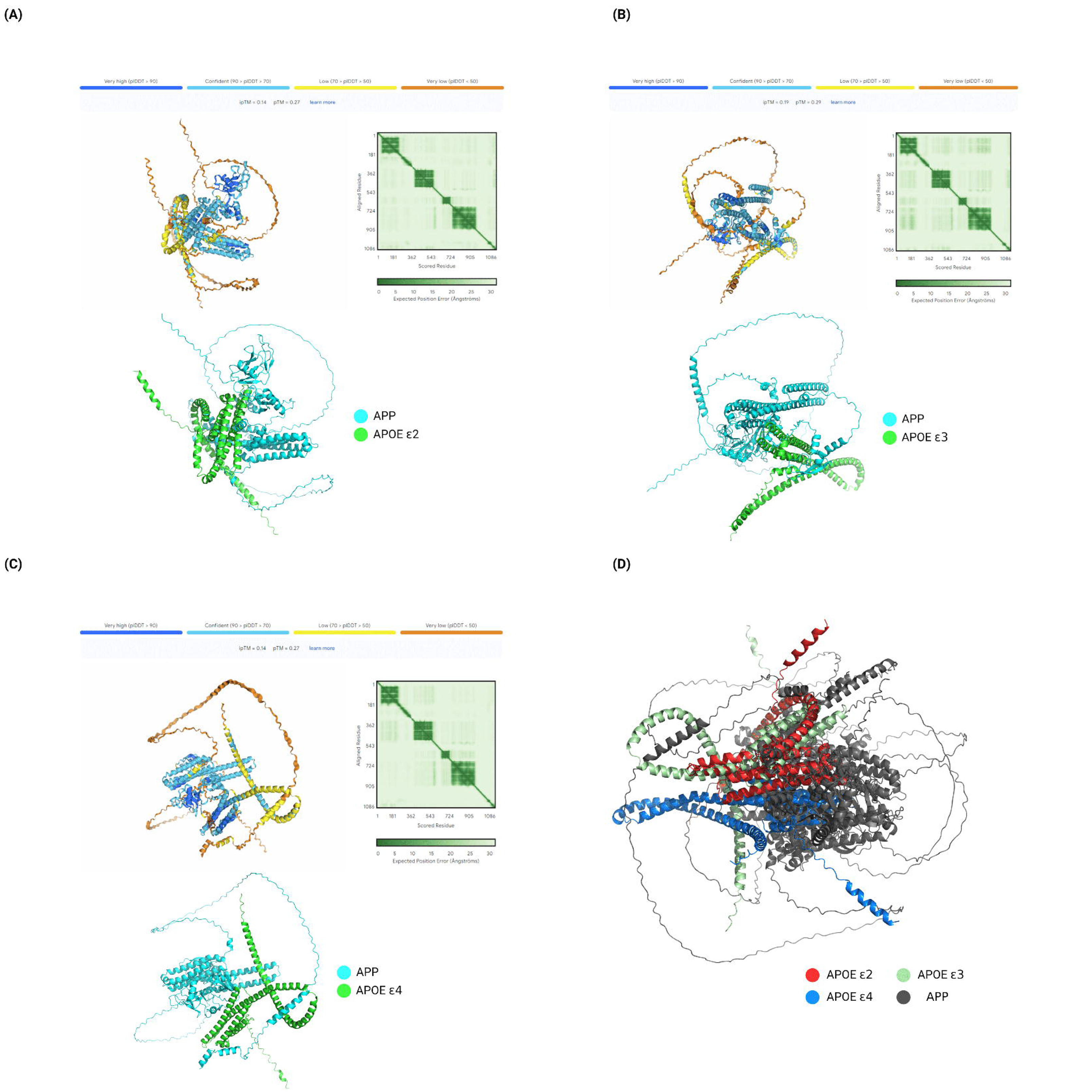
Structural prediction of APOE allele-APP complexes reveals isoform-specific binding modes associated with Alzheimer’s disease pathogenesis. (A) AlphaFold3-predicted structure of the APOE ε2-APP complex, with APOE ε2 shown in cyan and APP in gray. Confidence scores are mapped onto the structure with coloring as described in Figure 2. (B) Structural model of the APOE ε3-APP complex displaying the intermediate binding conformation between protective and pathogenic variants. (C) APOE ε4-APP complex structure showing the pathogenic binding mode characterized by altered interface orientation. (D) Superposition of all three APOE-APP complexes aligned by APP structure, revealing allele-specific differences in binding poses and interaction interfaces. The distinct conformational arrangements, particularly between ε2 (protective) and ε4 (pathogenic) variants, suggest that amino acid substitutions at positions 112 and 158 induce alterations in protein-protein recognition that may underlie their differential contributions to Alzheimer’s disease risk. These structural variations in binding geometry and interface composition represent potential targets for allele-selective therapeutic intervention.

For APOE-VLDLR interactions, structural predictions achieved pTM scores of 0.51, 0.54, and 0.51 for ε2, ε3, and ε4 variants, respectively (**Supplemental Fig.4**). Structural alignment revealed allele-specific variations in binding orientations. SASA analysis demonstrated surface area changes in 36 residues for ε2-VLDLR (mean: 0.402, variance: 0.100, SD: 0.317), 18 residues for ε3-VLDLR (mean: 0.320, variance: 0.034, SD: 0.185), and 44 residues for ε4-VLDLR (mean: 0.338, variance: 0.087, SD: 0.295) (**Supplemental Fig.3B & Supplemental Table 2**). This analysis identified 15, 8, and 15 interacting residues for the ε2, ε3, and ε4 complexes, respectively.

The APOE-ABCA1 complex predictions showed relatively higher confidence with pTM scores of 0.68, 0.65, and 0.68 for ε2, ε3, and ε4, respectively (**Supplemental Fig.5**). SASA analysis revealed 26 altered residues in ε2-ABCA1 (mean: 0.281, variance: 0.058, SD: 0.240), 32 residues in ε3-ABCA1 (mean: 0.216, variance: 0.026, SD: 0.161), and 10 residues in ε4-ABCA1 (mean: 0.273, variance: 0.027, SD: 0.165) (**Supplemental Fig.3C & Supplemental Table 3**). Notably, only the ε2-ABCA1 complex exhibited identifiable interaction residues (n=2), while no direct interaction residues were detected in the ε3 and ε4 complexes, suggesting potential differences in binding mechanisms.

Structural predictions for APOE-TREM2 complexes yielded lower confidence scores (pTM: 0.36, 0.36, and 0.34 for ε2, ε3, and ε4, respectively) (**Supplemental Fig.6**). Despite lower confidence, SASA analysis revealed consistent patterns with 22 altered residues in ε2-TREM2 (mean: 0.259, variance: 0.047, SD: 0.217), 23 residues in ε3-TREM2 (mean: 0.290, variance: 0.038, SD: 0.195), and 20 residues in ε4-TREM2 (mean: 0.236, variance: 0.038, SD: 0.195) (**Supplemental Fig.3D & Supplemental Table 4**). Eight, eight, and seven interacting residues were identified for the ε2, ε3, and ε4 complexes, respectively.

The APOE-APP complexes showed the lowest prediction confidence (pTM: 0.27, 0.29, and 0.27 for ε2, ε3, and ε4, respectively) (**Figure 3**). However, SASA analysis revealed substantial interface differences with 47 altered residues in ε2-APP (mean: 0.281, variance: 0.058, SD: 0.240), 50 residues in ε3-APP (mean: 0.359, variance: 0.088, SD: 0.296), and 35 residues in ε4-APP (mean: 0.299, variance: 0.049, SD: 0.220) (**Supplemental Fig.3E & Table 1**). The analysis identified 14, 16, and 14 interacting residues for the ε2, ε3, and ε4 complexes, respectively.

**Table 1.** Identification of allele-specific protein-protein interaction residues in APOE-APP complexes through structural analysis. (A) Interaction residues identified in the APOE ε2-APP complex using combined SASA analysis and PyMOL visualization. (B) Interaction residues mapped in the APOE ε3-APP complex interface. (C) Interaction residues characterized in the APOE ε4-APP complex revealing pathogenic-specific contact points. Residues are listed with their positions, amino acid identities, ΔSASA values (Å²), and inter-residue distances (Å) for contacts <5 Å. Bold entries indicate residues unique to specific APOE isoforms that may contribute to differential APP binding and Alzheimer’s disease pathogenesis.

| Analysis Tool | APOE ε2 | | | APP | |
| --- | --- | --- | --- | --- | --- |
|  | Residue | Amino Acid | Domain | Residue | Amino Acid |
| **SASA** | 30 | P | - | 718 | I |
| **SASA** | 33 | R | - | 722 | M |
| **SASA** | 90 | K | - | 769 | Q |
| **SASA, PyMOL** | 110 | R | - | 738 | V |
| **SASA, PyMOL** | 113 | K | - | 746 | E |
| **SASA** | 120 | A | - | 762 | Y |
| **SASA, PyMOL** | 121 | R | - | 745 | E |
| **SASA, PyMOL** | 127 | E | - | 759 | N |
| **SASA, PyMOL** | 127 | E | - | 761 | T |
| **SASA, PyMOL** | 168 | R | LDL and other lipoprotein receptors binding | 582 | S |
| **SASA, PyMOL** | 263 | E | Lipid-binding and lipoprotein association | 425 | K |
| **SASA** | 264 | Q | Lipid-binding and lipoprotein association | 704 | G |
| **SASA** | 267 | Q | Lipid-binding and lipoprotein association<br>Homooligomerization | 428 | K |
| **SASA, PyMOL** | 270 | L | Lipid-binding and lipoprotein association<br>Homooligomerization | 429 | K |
| **SASA, PyMOL** | 271 | Q | Lipid-binding and lipoprotein association<br>Homooligomerization | 429 | K |
| **SASA, PyMOL** | 278 | R | Lipid-binding and lipoprotein association<br>Homooligomerization<br>Specificity for association with VLDL | 437 | E |

(B)
| Analysis Tool | APOE ε3 | | | APP | |
| --- | --- | --- | --- | --- | --- |
|  | Residue | Amino Acid | Domain | Residue | Amino Acid |
| SASA | 46 | L | - | 332 | D |
| SASA | 50 | R | - | 336 | Y |
| SASA, PyMOL | 53 | D | - | 306 | R |
| SASA | 57 | W | - | 324 | C |
| SASA | 77 | E | - | 328 | R |
| SASA | 84 | E | - | 336 | Y |
| SASA | 110 | R | - | 767 | Q |
| SASA, PyMOL | 121 | R | - | 156 | E |
| SASA, PyMOL | 135 | Q | - | 529 | Q |
| SASA, PyMOL | 142 | A | - | 469 | R |
| SASA, PyMOL | 146 | Q | - | 544 | S |
| SASA | 150 | E | - | 540 | E |
| SASA, PyMOL | 154 | R | - | 539 | E |
| SASA, PyMOL | 164 | K | LDL and other lipoprotein receptors binding | 145 | E |
| SASA | 165 | R | LDL and other lipoprotein receptors binding | 152 | T |
| SASA, PyMOL | 168 | R | LDL and other lipoprotein receptors binding | 149 | H |

(C)
| Analysis Tool | APOE ε4 | | | APP | |
| --- | --- | --- | --- | --- | --- |
|  | Residue | Amino Acid | Domain | Residue | Amino Acid |
| SASA | 72 | S | - | 757 | Y |
| SASA | 76 | Q | - | 752 | M |
| SASA | 77 | E | - | 748 | H |
| SASA, PyMOL | 110 | R | - | 139 | E |
| SASA | 113 | K | - | 122 | F |
| SASA, PyMOL | 137 | R | - | 757 | Y |
| SASA | 141 | Q | - | 761 | T |
| SASA, PyMOL | 161 | K | LDL and other lipoprotein receptors binding | 200 | D |
| SASA | 172 | D | - | 129 | V |
| SASA, PyMOL | 175 | K | - | 131 | D |
| SASA | 176 | R | - | 127 | L |
| SASA | 207 | R | - | 706 | M |
| SASA | 214 | G | Lipid-binding and lipoprotein association | 705 | L |

### Domain-Specific Interaction Analysis and Target Selection

To understand the functional implications of identified interaction sites, we mapped them to known APOE functional domains: the LDL and lipoprotein receptor-binding region (residues 158-168), the lipid-binding and lipoprotein association domain (210-290), the homo-oligomerization region (266-317), and the VLDL-specific association domain (278-290) [70].

Analysis of domain-specific interactions revealed that most interaction sites with LDLR, VLDLR, and TREM2 localized to the LDL and lipoprotein receptor-binding region. However, these interactions showed minimal allele-specific variations in the actual interacting residues despite differences in overall binding poses. The ABCA1 interaction sites fell outside established functional domains, limiting their relevance for allele-specific targeting.

Significantly, the APP interaction analysis revealed distinct allele-specific differences that correlated with their opposing roles in Alzheimer’s disease. All identified APP interaction sites mapped to the LDL and lipoprotein receptor-binding domain, with a critical distinction: APOE ε2 engaged through R168, while APOE ε4 utilized R161. Although both residues reside within the same functional domain, their differential engagement suggests that amino acid variations at positions 112 and 158 induce subtle but functionally significant changes in APP interaction patterns.

Based on these comprehensive analyses, we selected the APOE-APP interaction as the primary target for inhibitor development. The allele-specific differences in APP binding, combined with APP’s central role in Aβ production and the opposing effects of ε2 (protective) and ε4 (risk-conferring) alleles, make this interaction an attractive therapeutic target for modulating Alzheimer’s disease pathogenesis.

### Target Selection and Peptide Inhibitor Design

The development of peptide-based inhibitors targeting protein-protein interaction interfaces represents a promising therapeutic strategy for modulating pathological interactions in Alzheimer’s disease [71]. Based on our structural analysis of the APOE ε4-APP complex, we designed peptide inhibitors to disrupt this critical interaction by targeting the identified hotspot residues on both binding partners.

Our inhibitor design strategy focused on two complementary approaches: (1) peptides targeting APOE ε4 interaction hotspots and (2) peptides targeting APP interaction sites. For APOE ε4-targeted inhibitors, we selected key residues identified from our interaction analysis: K161, D172, K175, and R176, which represent critical contact points in the APOE ε4-APP interface. For APP-targeted inhibitors, we focused on residues that directly interact with APOE ε4: L127 (interacts with APOE ε4 R176), V129 (interacts with APOE ε4 D172), and D131 (interacts with APOE ε4 K175).

### Computational Peptide Generation

Peptide backbone scaffolds were generated using RFdiffusion, with sequence lengths ranging from 6 to 16 amino acids. This length range was selected to balance multiple factors: the minimum of 6 residues ensures peptide-like properties and secondary structure formation, while the maximum of 16 residues maintains synthetic accessibility via standard solid-phase peptide synthesis and allows coverage of all targeted interaction sites. The backbone generation process incorporated spatial constraints to ensure optimal positioning relative to the target hotspots. This approach leverages artificial intelligence-based sequence design to identify amino acid combinations that stabilize the desired backbone conformations while maintaining favorable interaction profiles with the target sites. The design process involved iterative optimization cycles, generating approximately 11,000 candidate sequences for each target.

### Peptide Library Curation

After removing redundant sequences and filtering based on predicted structural stability, our final peptide libraries comprised 10,609 unique APOE ε4-targeting peptides and 10,661 APP-targeting peptides (**Supplemental Tables 5, 6**). These libraries represent a diverse collection of potential inhibitors with varying sequence compositions, predicted binding modes, and physicochemical properties. The large library size ensures comprehensive coverage of the chemical space around the target interaction sites, increasing the probability of identifying high-affinity inhibitors with favorable drug-like properties.

### Blood-Brain Barrier Permeability Assessment

Of the 10,609 APOE ε4-targeting peptides, 70 (0.66%) were predicted to possess BBB-penetrant properties. Similarly, 87 of 10,661 APP-targeting peptides (0.82%) demonstrated predicted BBB permeability. (**Supplemental Tables 5, 6**) While these represent relatively small fractions of the initial libraries, they provide focused sets of drug-like candidates that satisfy both target-binding and CNS-accessibility criteria.

### Molecular Docking Analysis and Binding Affinity Evaluation

Active residues for docking were defined based on our previously identified hotspots: K161, D172, K175, and R176 for APOE ε4, and L127, V129, D131, and D200 for APP. Surrounding residues were designated as passive residues to allow flexibility in binding mode exploration while maintaining focus on the critical interaction interfaces [72] (Supplemental Tables 7-10). A critical requirement for therapeutic efficacy is selectivity for the pathogenic APOE ε4 variant over the neutral ε3 and protective ε2 isoforms. To evaluate allele specificity, we performed comparative docking simulations of APOE-targeting peptides against all three APOE variants (**Table 2A**). This analysis identified nine peptides exhibiting preferential binding to APOE ε4, with average HADDOCK scores of approximately −41 kcal/mol for ε4 complexes compared to significantly weaker binding to ε2 and ε3 variants.

**Table 2.**
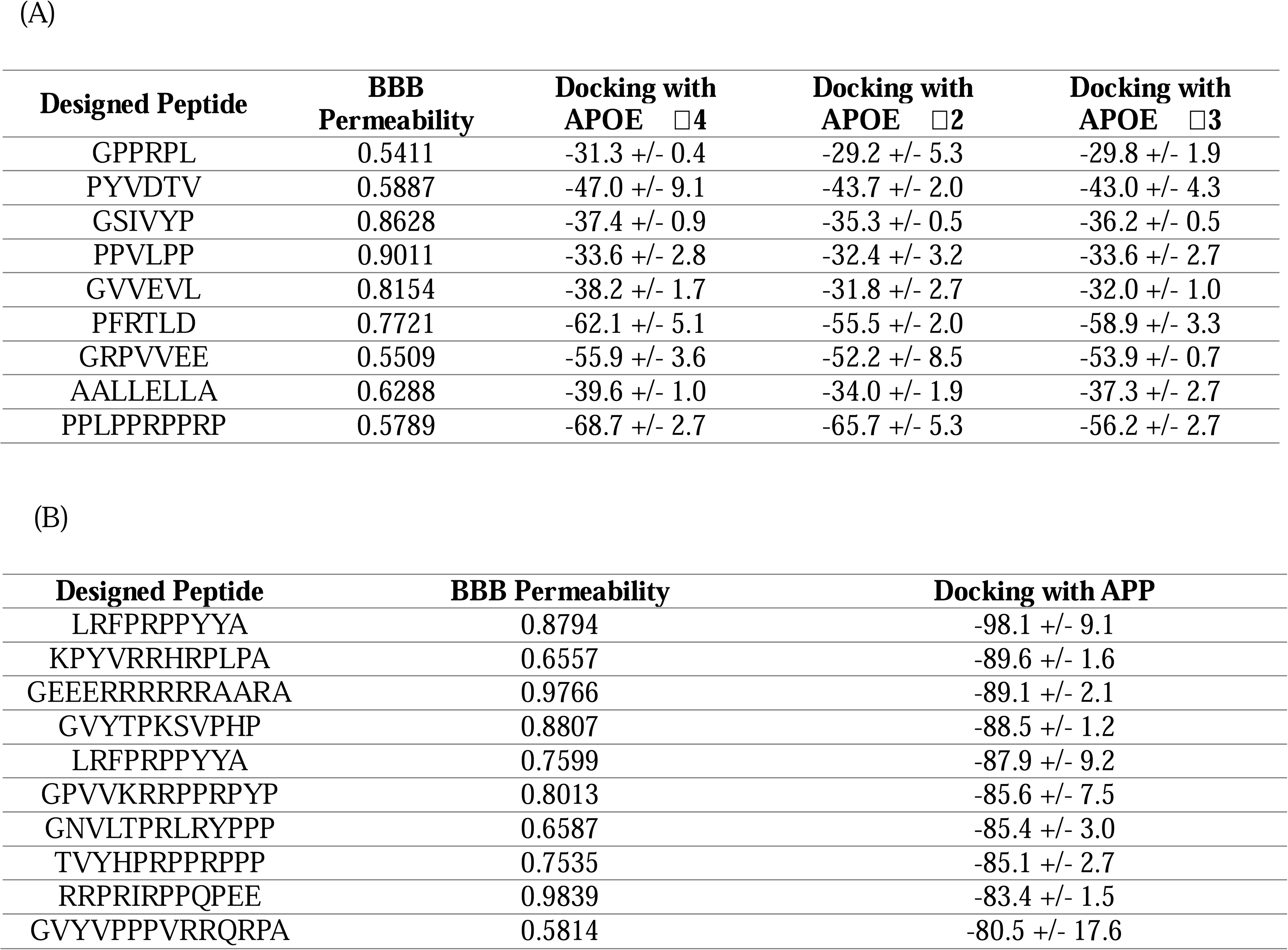
Lead peptide inhibitors with predicted blood-brain barrier permeability for targeting pathological APOE ε4-APP interactions. (A) Top nine APOE ε4-selective peptide inhibitors ranked by binding specificity, docking scores, and BBB permeability predictions. (B) Top ten APP-targeting peptides with corresponding binding affinities and BBB permeability assessments. All peptides were evaluated using DeepB3P3 for BBB penetration probability, HADDOCK for binding energy calculations, and structural stability metrics (RMSF, Rg, DCCM). Values represent means ± standard deviations where applicable.

For APP-targeting peptides, the top 10 candidates based on docking scores were selected for detailed analysis (**Table 2B**). These peptides demonstrated stronger predicted binding affinities with average HADDOCK scores of −64 kcal/mol, suggesting potentially more favorable thermodynamic interactions compared to APOE-targeting peptides. However, binding affinity alone does not determine therapeutic potential, necessitating comprehensive structural stability assessments.

### Structural Stability and Dynamic Analysis

To evaluate the structural integrity and dynamic behavior of peptide-target complexes, we performed molecular dynamics-based analyses using three complementary metrics: Root Mean Square Fluctuation (RMSF) for residue-level flexibility, Radius of Gyration (Rg) for overall compactness, and Dynamic Cross-Correlation Matrix (DCCM) for correlated motions between residues.

RMSF analysis revealed distinct stability profiles between the two peptide classes. APOE ε4-specific binders exhibited average RMSF values of approximately 21 Å, indicating relatively rigid conformations upon binding. In contrast, APP binders showed higher average RMSF values of approximately 31 Å, suggesting greater conformational flexibility (**Table 3A, Supplemental Table 11**). Lower RMSF values typically correlate with more stable protein-peptide interactions, favoring the APOE ε4-targeting peptides in terms of binding stability.

**Table 3.**
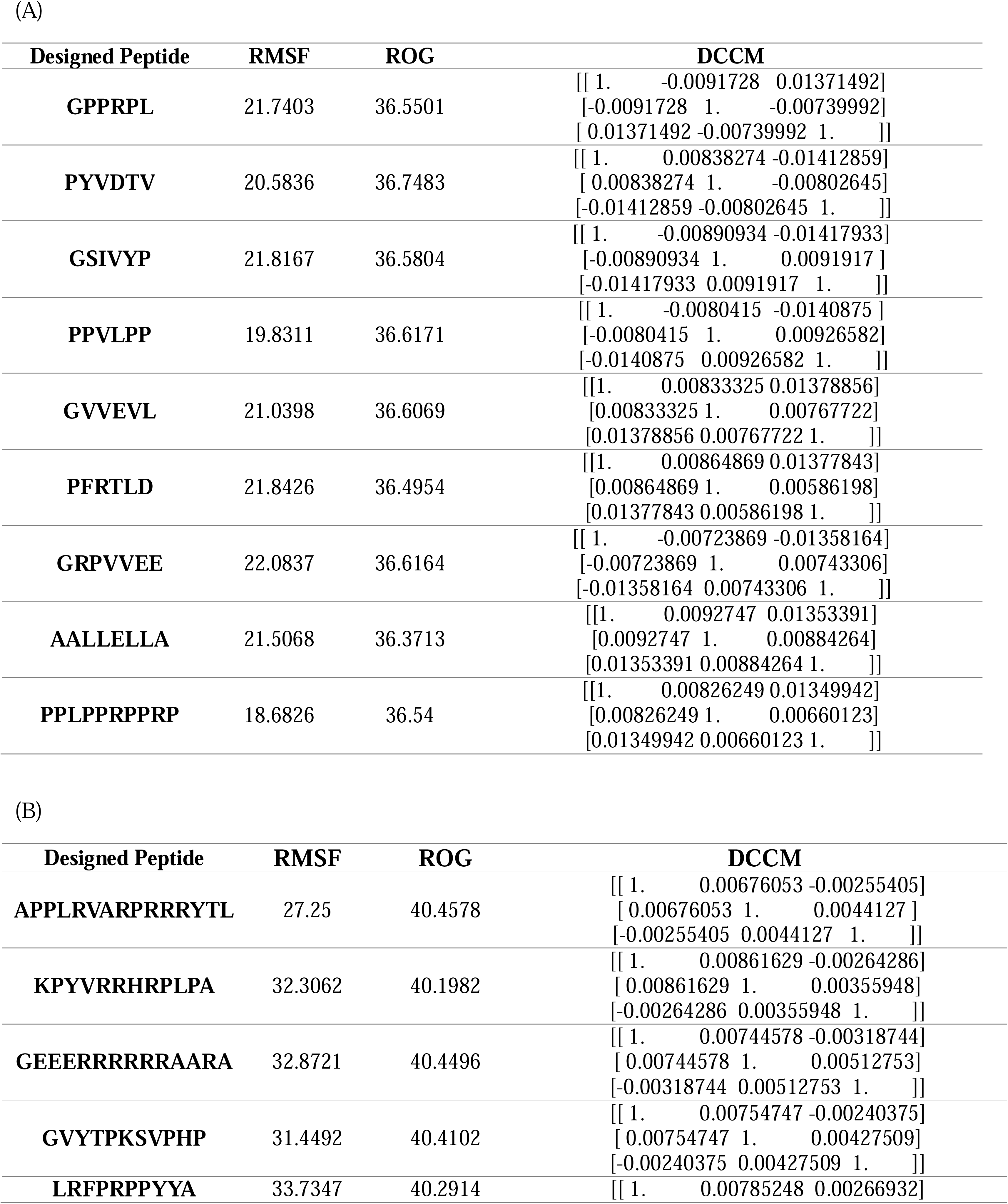
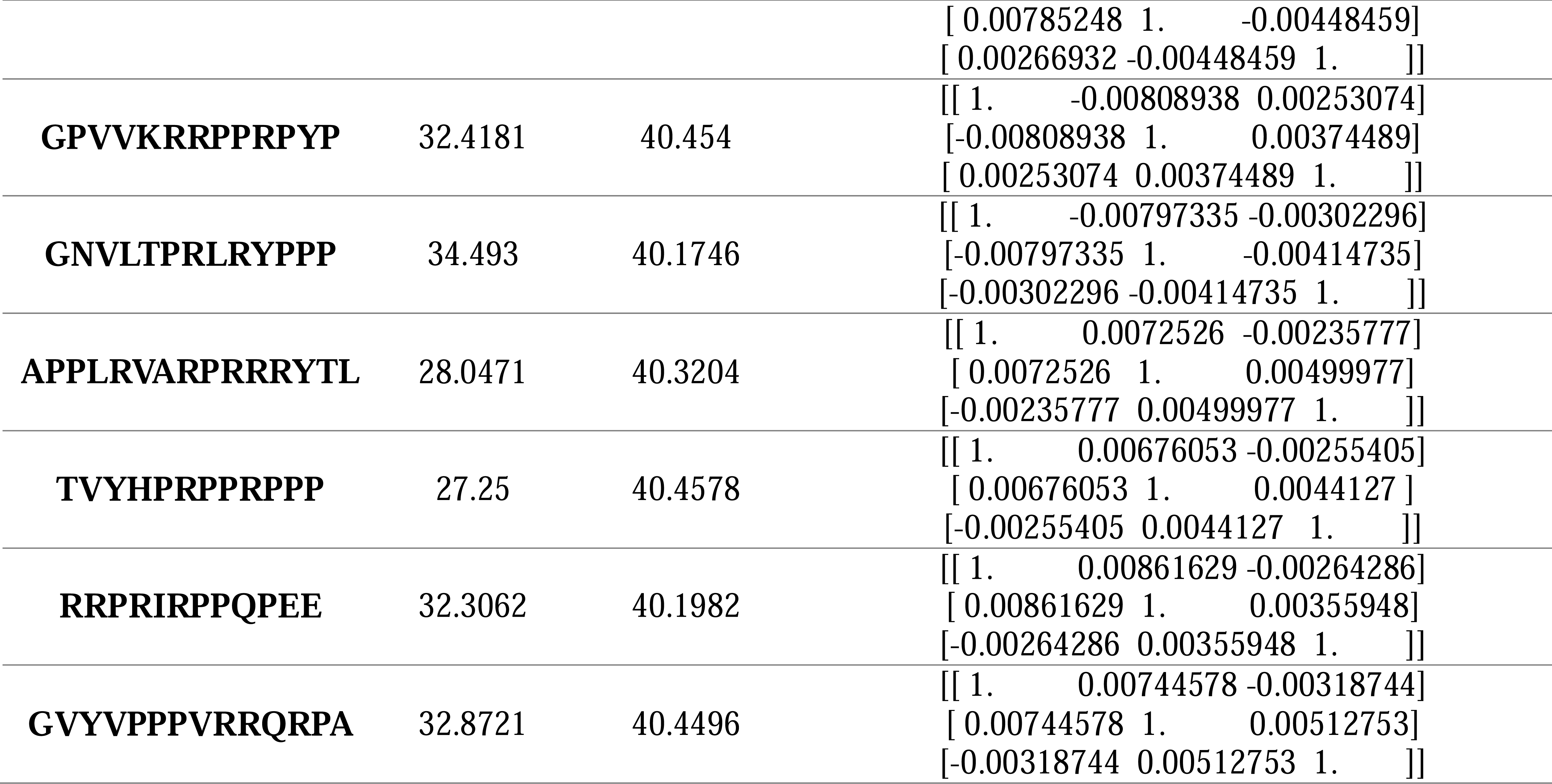
Structural stability and dynamic characterization of lead peptide inhibitors. (A) Molecular dynamics-based stability parameters for nine APOE ε4-selective peptide inhibitors, including root mean square fluctuation (RMSF), radius of gyration (Rg), and dynamic cross-correlation matrix (DCCM) analysis. (B) Comparative stability assessment of top ten APP-targeting peptides using identical structural metrics. Values indicate conformational flexibility, compactness, and correlated motions that determine binding stability and specificity. Peptides exhibiting RMSF <25 Å, appropriate Rg for their sequence length, and stable DCCM patterns are considered optimal candidates for further development.

Radius of gyration measurements provided insights into the overall structural compactness of bound peptides. APOE ε4-specific binders displayed average Rg values of approximately 36 Å, while APP binders showed more compact structures with average Rg values of 31 Å (**Table 3B, Supplemental Table 12**) However, this difference primarily reflects the varying peptide lengths rather than intrinsic folding propensities. When normalized for sequence length, APOE ε4-targeting peptides demonstrated more extended conformations, potentially allowing better surface complementarity with their target.

DCCM analysis evaluated the internal dynamic coupling within peptide structures, providing insights into their conformational stability during binding. Among APOE ε4-specific binders, four peptides (ε4_binder_6-918, ε4_binder_6-999, ε4_binder_8-144, and ε4_binder_10-142) exhibited stable correlated motions indicative of well-defined binding conformations (**Table 3A, B**). Conversely, none of the APP-targeting peptides displayed similar stability in their correlated motions, suggesting more dynamic and potentially less specific binding modes.

### Lead Candidates Selection

Despite APP-targeting peptides showing more favorable docking scores, the comprehensive structural analysis revealed superior stability characteristics for APOE ε4-specific binders. The combination of lower conformational flexibility, appropriate structural extension for target engagement, and stable correlated motions positions APOE ε4-targeting peptides as more promising candidates for interaction inhibition.

Based on the comprehensive evaluation encompassing BBB permeability, allele-specific binding affinity, and structural stability analyses, we identified nine APOE ε4-specific peptide binders as lead candidates. These nine peptides (GPPRPL, PYVDTV, GSIVYP, PPVLPP, GVVEVL, PFRTLD, GRPVVEE, AALLELLA, and PPLPPRPPRP) demonstrated optimal characteristics across all assessment criteria: (1) predicted BBB permeability enabling CNS delivery, (2) selective binding to APOE ε4 over ε2 and ε3 isoforms with favorable docking scores, (3) superior structural stability with low RMSF values, and (4) stable correlated motions indicating well-defined binding conformations.

## Discussion

This study leveraged artificial intelligence-driven computational approaches to elucidate the molecular basis of APOE ε4-mediated risk in Alzheimer’s disease and develop targeted therapeutic interventions. Through comprehensive structural analysis of APOE allele-specific protein interactions, we identified critical differences in the APOE ε4-APP interaction interface that distinguish the pathogenic ε4 variant from the protective ε2 and neutral ε3 isoforms.

Our structural investigations revealed that while APOE alleles maintain highly conserved overall architectures, subtle variations at the interaction interfaces manifest as functionally significant differences in protein-protein interactions. The identification of allele-specific interaction patterns, particularly the differential engagement of R161 in APOE ε4 versus R168 in APOE ε2 within the APP binding interface, provides molecular insights into the opposing roles these variants play in disease pathogenesis. These findings align with and extend previous biochemical studies suggesting that APOE ε4 compromises Aβ clearance mechanisms while promoting aggregation pathways

The rational design of peptide inhibitors targeting the APOE ε4-APP interaction represents a novel therapeutic strategy that directly addresses the molecular mechanisms underlying genetic risk. By employing state-of-the-art computational tools including RFdiffusion for backbone generation and ProteinMPNN for sequence optimization, we generated extensive peptide libraries that were systematically evaluated for drug-like properties. The identification of nine lead candidates that combine BBB permeability, allele selectivity, and structural stability demonstrates the power of integrated computational pipelines in drug discovery.

Notably, our multi-parametric evaluation approach revealed important considerations for peptide drug design. While APP-targeting peptides showed superior binding affinities in docking simulations, APOE ε4-targeting peptides demonstrated more favorable structural stability profiles, highlighting the importance of comprehensive assessment beyond simple binding energy calculations. The superior performance of APOE ε4-specific binders in RMSF and DCCM analyses suggests that targeting the more structured receptor-binding domain of APOE may provide more stable and specific therapeutic interactions compared to targeting the inherently flexible APP structure.

The integration of multiple computational methodologies—from AlphaFold3 structure prediction to molecular dynamics-based stability assessments—exemplifies how modern AI algorithms can accelerate the drug discovery process. The DeepB3P3 algorithm’s identification of BBB-permeable candidates addresses a critical challenge in CNS drug development, potentially reducing the high attrition rates associated with neurotherapeutic development. Furthermore, the allele-specific targeting achieved through comparative docking simulations offers the possibility of precision medicine approaches that selectively modulate pathogenic interactions while preserving normal protein functions.

Several limitations warrant consideration. First, the moderate pTM scores obtained for some protein complexes reflect the current limitations in predicting large, flexible protein-protein interactions, particularly for proteins lacking extensive experimental structural data. Second, while our computational pipeline provides valuable insights, the predicted binding modes and affinities require experimental validation through biophysical characterization and cellular assays. Third, the dynamic nature of protein-protein interactions in cellular environments, including effects of post-translational modifications and membrane associations, cannot be fully captured by current computational methods.

Future studies should focus on experimental validation of the identified lead peptides through surface plasmon resonance or isothermal titration calorimetry to confirm binding affinities and specificities. Cell-based assays examining the ability of these peptides to disrupt APOE ε4-APP interactions and modulate Aβ metabolism will be essential for establishing therapeutic relevance. Additionally, optimization of peptide stability through chemical modifications such as cyclization or D-amino acid incorporation may enhance their therapeutic potential.

The broader implications of this work extend beyond Alzheimer’s disease. The computational framework established here—combining structure prediction, interaction analysis, and rational inhibitor design—can be applied to other protein misfolding disorders and diseases driven by pathological protein-protein interactions. As AI algorithms continue to improve in accuracy and scope, we anticipate that such structure-based approaches will become increasingly valuable for understanding disease mechanisms and developing targeted therapeutics.

In conclusion, this study demonstrates the potential of AI-driven structural biology to bridge the gap between genetic risk factors and therapeutic intervention. By revealing the structural basis of APOE ε4 pathogenicity and designing selective inhibitors of pathological interactions, we provide a foundation for developing precision therapeutics for Alzheimer’s disease. The successful integration of multiple computational approaches highlights the transformative potential of artificial intelligence in modern drug discovery, while emphasizing the continued importance of experimental validation in translating computational insights into clinical applications.

## Supporting information

Manuscript

## Acknowledgement

This work was supported by the National Research Foundation of Korea (NRF) grant funded by the Korean government (MSIT) (RS-2023-00209456) and the Korea Basic Science Institute (National Research Facilities and Equipment Center) grant funded by the Korean government (MSIT) (RS-2024-00402298). We acknowledge the use of Claude Opus 4.1 solely for linguistic refinement and grammatical corrections in manuscript preparation. All scientific content, data analysis, and intellectual contributions presented herein were developed independently by the authors without the use of generative AI tools. Figure 1 was created with BioRender.com (accessed 1 July 2025).

## Author Contributions

J.J., E.H, and J.P: computational analysis, peptide design, molecular docking simulations, data analysis, writing-original draft. A.S.: Conceptualization, project administration, resources, supervision, writing-review & editing. H.K.: Conceptualization, methodology, project administration, funding acquisition, supervision, writing-review & editing. All authors have read and approved the final manuscript.

---

**Supplemental Fig. 1. AlphaFold3 structural predictions of individual APOE allele variants.**

Predicted three-dimensional structures of APOE isoforms generated using AlphaFold3. (A) APOE ε2 (Cys112/Cys158) structure with pTM score of 0.51. (B) APOE ε3 (Cys112/Arg158) structure with pTM score of 0.52. (C) APOE ε4 (Arg112/Arg158) structure with pTM score of 0.51. Structures are colored according to pLDDT confidence scores: blue (very high confidence, pLDDT >90), cyan (high confidence, pLDDT 70-90), yellow (moderate confidence, pLDDT 50-70), and orange (low confidence, pLDDT <50). The moderate pTM scores reflect the limited availability of experimentally determined full-length APOE structures in existing databases. (D) Structural alignment of all three APOE allele structure.

**Supplemental Fig. 2. Structural prediction of APOE allele-LDLR complexes.**

AlphaFold3-predicted structures of APOE variants in complex with low-density lipoprotein receptor (LDLR). (A) APOE ε2-LDLR complex structure (pTM = 0.43). (B) APOE ε3-LDLR complex structure (pTM = 0.45). (C) APOE ε4-LDLR complex structure (pTM = 0.48). (D) Structural alignment of all three APOE-LDLR complexes based on LDLR structure, revealing distinct binding poses across alleles. APOE proteins are shown in different colors (ε2 in cyan, ε3 in green, ε4 in magenta) while LDLR is displayed in gray. The differential binding orientations suggest allele-specific variations in receptor recognition despite the conserved LDL receptor-binding domain.

**Supplemental Fig. 3. Solvent-accessible surface area (SASA) analysis reveals allele-specific differences in APOE protein-protein interaction interfaces.**

(A) Comparative SASA analysis of APOE ε2, ε3, and ε4 complexes with LDLR. Residues exhibiting significant changes in solvent accessibility upon complex formation are highlighted, revealing allele-specific variations in the binding interface. (B) SASA-based identification of interaction residues in APOE allele-VLDLR complexes, demonstrating differential surface area changes across isoforms. (C) Interface analysis of APOE variants bound to ABCA1, showing distinct patterns of surface accessibility changes with notable absence of direct interaction residues in ε3 and ε4 complexes. (D) SASA profiling of APOE allele-TREM2 interactions, identifying conserved and variant-specific interface residues. (E) Differential SASA analysis of APOE-APP complexes across three isoforms, revealing allele-specific interaction patterns that correlate with their opposing roles in Alzheimer’s disease pathogenesis. Bar graphs represent the magnitude of SASA changes (Δ SASA in Å²) for individual residues, with statistical significance indicated for residues showing >20% change in surface accessibility upon binding.

**Supplemental Fig. 4. Structural prediction of APOE allele-VLDLR complexes.**

AlphaFold3-generated structures of APOE isoforms bound to very low-density lipoprotein receptor (VLDLR). (A) APOE ε2-VLDLR complex with pTM score of 0.51. (B) APOE ε3-VLDLR complex with pTM score of 0.54, showing the highest confidence among the three variants. (C) APOE ε4-VLDLR complex with pTM score of 0.51. (D) Superposition of all three complexes aligned by VLDLR structure, demonstrating allele-specific variations in binding orientations. The structural alignment reveals differences in binding poses that may contribute to the differential association of APOE ε4 with VLDL-rich particles compared to APOE ε2’s preference for HDL-rich particles.

**Supplemental Fig. 5. Structural prediction of APOE allele-ABCA1 complexes.**

Predicted structures of APOE variants in complex with ATP-binding cassette transporter A1 (ABCA1), a critical regulator of APOE lipidation. (A) APOE ε2-ABCA1 complex structure (pTM = 0.68). (B) APOE ε3-ABCA1 complex structure (pTM = 0.65). (C) APOE ε4-ABCA1 complex structure (pTM = 0.68). (D) Structural comparison of all three APOE-ABCA1 complexes aligned by ABCA1 transmembrane domains. The relatively high pTM scores indicate robust prediction confidence. ABCA1 is shown as a gray surface representation with transmembrane helices highlighted. Despite similar confidence scores, subsequent SASA analysis revealed marked differences in interaction patterns, with only the ε2 variant showing identifiable direct interaction residues, suggesting distinct binding mechanisms that may explain APOE ε4’s reduced lipidation efficiency.

**Supplemental Fig. 6. Structural prediction of APOE allele-TREM2 complexes.**

AlphaFold3 predictions of APOE isoforms bound to triggering receptor expressed on myeloid cells 2 (TREM2), a microglial receptor implicated in Alzheimer’s disease pathology. (A) APOE ε2-TREM2 complex (pTM = 0.36). APOE ε3-TREM2 complex (pTM = 0.36). (C) APOE ε4-TREM2 complex (pTM = 0.34). (D) Overlay of all three APOE-TREM2 complexes showing variations in binding geometry. The lower pTM scores reflect the challenging nature of predicting this interaction, potentially due to the lipidation-dependent binding mechanism. TREM2 is displayed in gray with its immunoglobulin-like domain highlighted. Despite lower confidence scores, consistent interaction patterns were identified, with APOE ε4 showing reduced binding that may contribute to impaired microglial function in Alzheimer’s disease.

