## Supplementary material for "*In Silico* Design of APOE ɛ4 Interaction Inhibitor Peptides for Alzheimer’s Disease": Manuscript: Supplemental Table 1.docx

**Table 1. List of PPIs from APOE Allele(ɛ2, ɛ3, ɛ4) - LDLR Complex**

(A) List of identified protein-protein interaction in APOE ɛ2 Allele-LDLR interactome by SASA analysis and PyMOL.

| Analysis Tool | APOE ɛ2 | | | LDLR | |
| --- | --- | --- | --- | --- | --- |
|  | Residue | Amino Acid | Domain | Residue | Amino Acid |
| SASA | 29 | E | - | 187 | L |
| SASA, PyMOL | 132 | R | - | 835 | E |
| SASA, PyMOL | 139 | E | - | 828 | Y |
| SASA | 146 | Q | - | 817 | N |
| SASA | 146 | Q | - | 819 | N |
| SASA, PyMOL | 154 | R | - | 823 | F |
| SASA | 158 | H | LDL and other lipoprotein receptors binding | 832 | T |
| SASA | 160 | R | LDL and other lipoprotein receptors binding | 170 | D |
| SASA, PyMOL | 164 | K | LDL and other lipoprotein receptors binding | 168 | D |
| SASA, PyMOL | 164 | K | LDL and other lipoprotein receptors binding | 172 | D |
| SASA, PyMOL | 165 | R | LDL and other lipoprotein receptors binding | 836 | V |

(B) List of identified protein-protein interaction in APOE ɛ3 Allele-LDLR interactome by SASA analysis and PyMOL.

| Analysis Tool | APOE ɛ3 | | | LDLR | |
| --- | --- | --- | --- | --- | --- |
|  | Residue | Amino Acid | Domain | Residue | Amino Acid |
| SASA, PyMOL | 106 | E | - | 850 | R |
| SASA, PyMOL | 113 | K | - | 835 | E |
| SASA | 114 | E | - | 847 | Y |
| SASA | 117 | A | - | 832 | T |
| SASA, PyMOL | 121 | R | - | 836 | V |
| SASA, PyMOL | 128 | D | - | 830 | K |
| SASA | 132 | R | - | 824 | D |
| SASA, PyMOL | 139 | E | - | 819 | N |
| SASA, PyMOL | 154 | R | - | 816 | N |
| SASA | 158 | H | LDL and other lipoprotein receptors binding | 821 | I |
| SASA, PyMOL | 160 | R | LDL and other lipoprotein receptors binding | 169 | N |
| SASA, PyMOL | 164 | K | LDL and other lipoprotein receptors binding | 170 | D |
| SASA, PyMOL | 164 | K | LDL and other lipoprotein receptors binding | 172 | D |
| SASA, PyMOL | 165 | R | LDL and other lipoprotein receptors binding | 833 | E |
| SASA | 168 | R | - | 833 | E |
| SASA | 176 | R | - | 838 | I |

(C) List of identified protein-protein interaction in APOE ɛ4 Allele-LDLR interactome by SASA analysis and PyMOL.

| Analysis Tool | APOE ɛ4 | | | LDLR | |
| --- | --- | --- | --- | --- | --- |
|  | Residue | Amino Acid | Domain | Residue | Amino Acid |
| SASA, PyMOL | 29 | E | - | 313 | K |
| SASA | 31 | E | - | 313 | K |
| SASA, PyMOL | 53 | D | - | 620 | K |
| SASA, PyMOL | 56 | R | - | 619 | S |
| SASA, PyMOL | 56 | R | - | 645 | A |
| SASA, PyMOL | 56 | R | - | 646 | H |
| SASA, PyMOL | 63 | E | - | 491 | K |
| SASA | 128 | D | - | 210 | W |
| SASA | 132 | R | - | 197 | E |
| SASA, PyMOL | 149 | E | - | 448 | R |
| SASA, PyMOL | 158 | H | LDL and other lipoprotein receptors binding | 197 | E |
| SASA, PyMOL | 160 | R | LDL and other lipoprotein receptors binding | 663 | E |
| SASA, PyMOL | 161 | K | LDL and other lipoprotein receptors binding | 217 | D |
| SASA, PyMOL | 165 | R | LDL and other lipoprotein receptors binding | 213 | D |
| SASA, PyMOL | 168 | R | LDL and other lipoprotein receptors binding | 215 | D |
