## Supplementary material for "*In Silico* Design of APOE ɛ4 Interaction Inhibitor Peptides for Alzheimer’s Disease": Manuscript: Supplemental Table 2.docx

**Table 2. List of PPIs from APOE Allele(ɛ2, ɛ3, ɛ4) - VLDLR Complex**

(A) List of identified protein-protein interaction in APOE ɛ2 Allele-VLDLR interactome by SASA analysis and PyMOL.

| Analysis Tool | APOE ɛ2 | | | VLDLR | |
| --- | --- | --- | --- | --- | --- |
|  | Residue | Amino Acid | Domain | Residue | Amino Acid |
| SASA, PyMOL | 34 | Q | - | 196 | H |
| SASA | 106 | E | - | 535 | K |
| SASA, PyMOL | 110 | R | - | 557 | E |
| SASA | 113 | K | - | 830 | M |
| SASA, PyMOL | 114 | E | - | 841 | T |
| SASA | 158 | H | LDL and other lipoprotein receptors binding | 253 | H |
| SASA, PyMOL | 160 | K | LDL and other lipoprotein receptors binding | 213 | D |
| SASA, PyMOL | 160 | K | LDL and other lipoprotein receptors binding | 215 | D |
| SASA, PyMOL | 161 | K | LDL and other lipoprotein receptors binding | 259 | D |
| SASA, PyMOL | 161 | K | LDL and other lipoprotein receptors binding | 263 | D |
| SASA, PyMOL | 165 | K | LDL and other lipoprotein receptors binding | 261 | D |
| SASA | 179 | V | - | 844 | D |
| SASA, PyMOL | 186 | E | - | 491 | K |

(B) List of identified protein-protein interaction in APOE ɛ3 Allele-VLDLR interactome by SASA analysis and PyMOL.

| Analysis Tool | APOE ɛ3 | | | VLDLR | |
| --- | --- | --- | --- | --- | --- |
|  | Residue | Amino Acid | Domain | Residue | Amino Acid |
| SASA, PyMOL | 113 | K | - | 74 | E |
| SASA, PyMOL | 121 | R | - | 851 | R |
| SASA | 157 | S | - | 728 | H |
| SASA, PyMOL | 161 | K | LDL and other lipoprotein receptors binding | 727 | D |
| SASA, PyMOL | 169 | D | - | 851 | R |
| SASA | 299 | E | Homooligomerization | 777 | P |
| SASA | 299 | E | Homooligomerization | 778 | G |
| SASA | 299 | E | Homooligomerization | 779 | G |

(C) List of identified protein-protein interaction in APOE ɛ4 Allele-VLDLR interactome by SASA analysis and PyMOL.

| Analysis Tool | APOE ɛ4 | | | VLDLR | |
| --- | --- | --- | --- | --- | --- |
|  | Residue | Amino Acid | Domain | Residue | Amino Acid |
| SASA, PyMOL | 29 | E | - | 313 | K |
| SASA, PyMOL | 53 | D | - | 620 | K |
| SASA, PyMOL | 56 | R | - | 646 | H |
| SASA, PyMOL | 56 | R | - | 645 | A |
| SASA, PyMOL | 56 | R | - | 619 | S |
| SASA, PyMOL | 63 | E | - | 491 | K |
| SASA, PyMOL | 149 | E | - | 448 | R |
| SASA, PyMOL | 158 | H | LDL and other lipoprotein receptors binding | 197 | E |
| SASA, PyMOL | 160 | R | LDL and other lipoprotein receptors binding | 663 | E |
| SASA, PyMOL | 161 | K | LDL and other lipoprotein receptors binding | 217 | D |
| SASA | 164 | K | LDL and other lipoprotein receptors binding | 664 | N |
| SASA, PyMOL | 165 | R | LDL and other lipoprotein receptors binding | 213 | D |
| SASA, PyMOL | 168 | R | LDL and other lipoprotein receptors binding | 215 | D |
| SASA | 189 | E | - | 298 | G |
| SASA | 264 | Q | Lipid-binding and lipoprotein association Specificity for association with VLDL | 839 | K |
