## Supplementary material for "*In Silico* Design of APOE ɛ4 Interaction Inhibitor Peptides for Alzheimer’s Disease": Manuscript: Supplemental Table 3.docx

**Table 4. List of PPIs from APOE Allele(ɛ2, ɛ3, ɛ4) – ABCA1 Complex**

(A) List of identified protein-protein interaction in APOE ɛ2 Allele-ABCA1 interactome by SASA analysis and PyMOL.

| Analysis Tool | APOE ɛ2 | | | ABCA1 | |
| --- | --- | --- | --- | --- | --- |
|  | Residue | Amino Acid | Domain | Residue | Amino Acid |
| SASA | 10 | R | - | 1371 | F |
| SASA | 113 | K | - | 1368 | V |
