## Supplementary material for "*In Silico* Design of APOE ɛ4 Interaction Inhibitor Peptides for Alzheimer’s Disease": Manuscript: Supplemental Table 4.docx

**Table 3. List of PPIs from APOE Allele(ɛ2, ɛ3, ɛ4) – TREM2 Complex**

(A) List of identified protein-protein interaction in APOE ɛ2 Allele-TERM2 interactome by SASA analysis and PyMOL.

| Analysis Tool | APOE ɛ2 | | | TREM2 | |
| --- | --- | --- | --- | --- | --- |
|  | Residue | Amino Acid | Domain | Residue | Amino Acid |
| SASA, PyMOL | 116 | Q |  | 87 | D |
| SASA | 120 | A |  | 33 | Q |
| SASA, PyMOL | 127 | E |  | 76 | R |
| SASA, PyMOL | 127 | E |  | 98 | R |
| SASA, PyMOL | 128 | D |  | 31 | S |
| SASA, PyMOL | 161 | K | LDL and other lipoprotein receptors binding | 131 | D |
| SASA, PyMOL | 161 | K | LDL and other lipoprotein receptors binding | 134 | D |
| SASA | 165 | R | LDL and other lipoprotein receptors binding | 31 | S |

(B) List of identified protein-protein interaction in APOE ɛ3 Allele- TERM2 interactome by SASA analysis and PyMOL.

| Analysis Tool | APOE ɛ3 | | | TREM2 | |
| --- | --- | --- | --- | --- | --- |
|  | Residue | Amino Acid | Domain | Residue | Amino Acid |
| SASA, PyMOL | 116 | Q |  | 87 | D |
| SASA, PyMOL | 127 | E |  | 76 | R |
| SASA, PyMOL | 127 | E |  | 98 | R |
| SASA, PyMOL | 128 | D |  | 31 | S |
| SASA, PyMOL | 128 | D |  | 98 | R |
| SASA, PyMOL | 161 | K | LDL and other lipoprotein receptors binding | 131 | D |
| SASA | 165 | R | LDL and other lipoprotein receptors binding | 31 | S |
| SASA | 278 | R | Lipid-binding and lipoprotein association Homooligomerization Specificity for association with VLDL | 44 | W |

(C) List of identified protein-protein interaction in APOE ɛ4 Allele- TERM2 interactome by SASA analysis and PyMOL.

| Analysis Tool | APOE ɛ4 | | | TREM2 | |
| --- | --- | --- | --- | --- | --- |
|  | Residue | Amino Acid | Domain | Residue | Amino Acid |
| SASA, PyMOL | 127 | E |  | 76 | R |
| SASA, PyMOL | 128 | D |  | 31 | S |
| SASA | 131 | G |  | 98 | R |
| SASA, PyMOL | 135 | Q |  | 98 | R |
| SASA, PyMOL | 165 | R | LDL and other lipoprotein receptors binding | 29 | G |
| SASA, PyMOL | 168 | R | LDL and other lipoprotein receptors binding | 131 | D |
| SASA | 271 | Q | Lipid-binding and lipoprotein association Homooligomerization | 44 | W |
