## Supplementary figures and images for "*In Silico* Design of APOE ɛ4 Interaction Inhibitor Peptides for Alzheimer’s Disease"

### Supplemental Fig.1.png

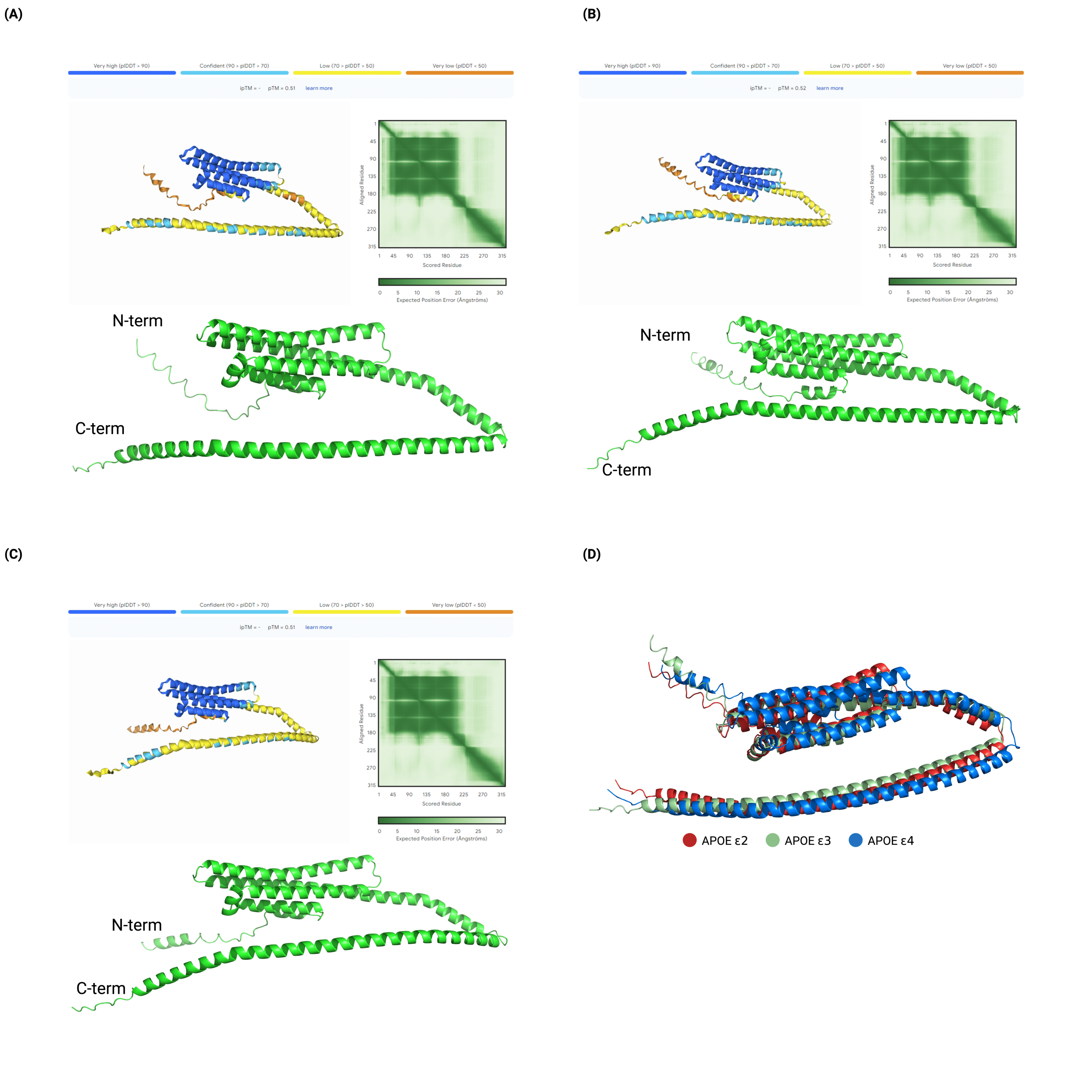

### Supplemental Fig.2.png

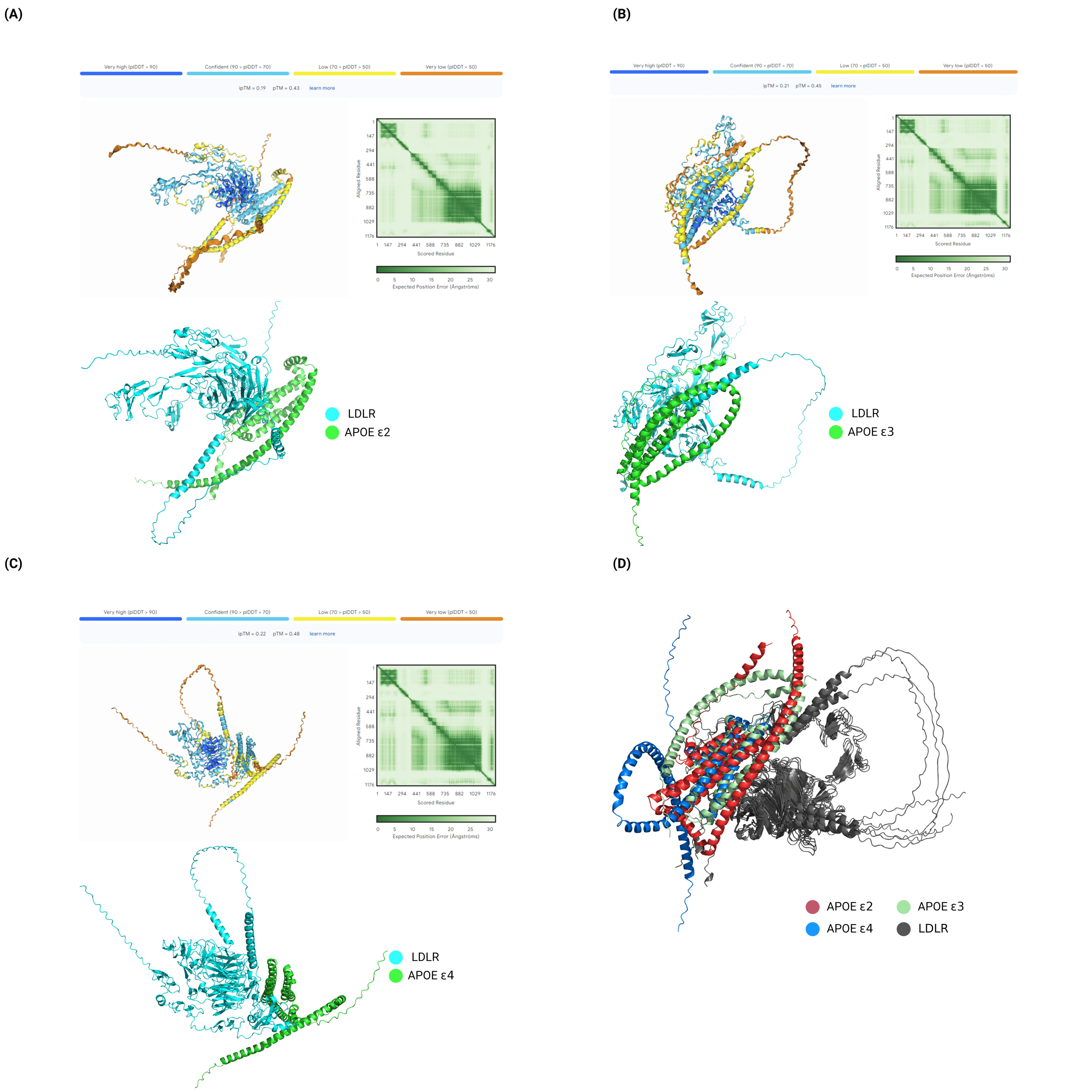

### Supplemental Fig.3.png

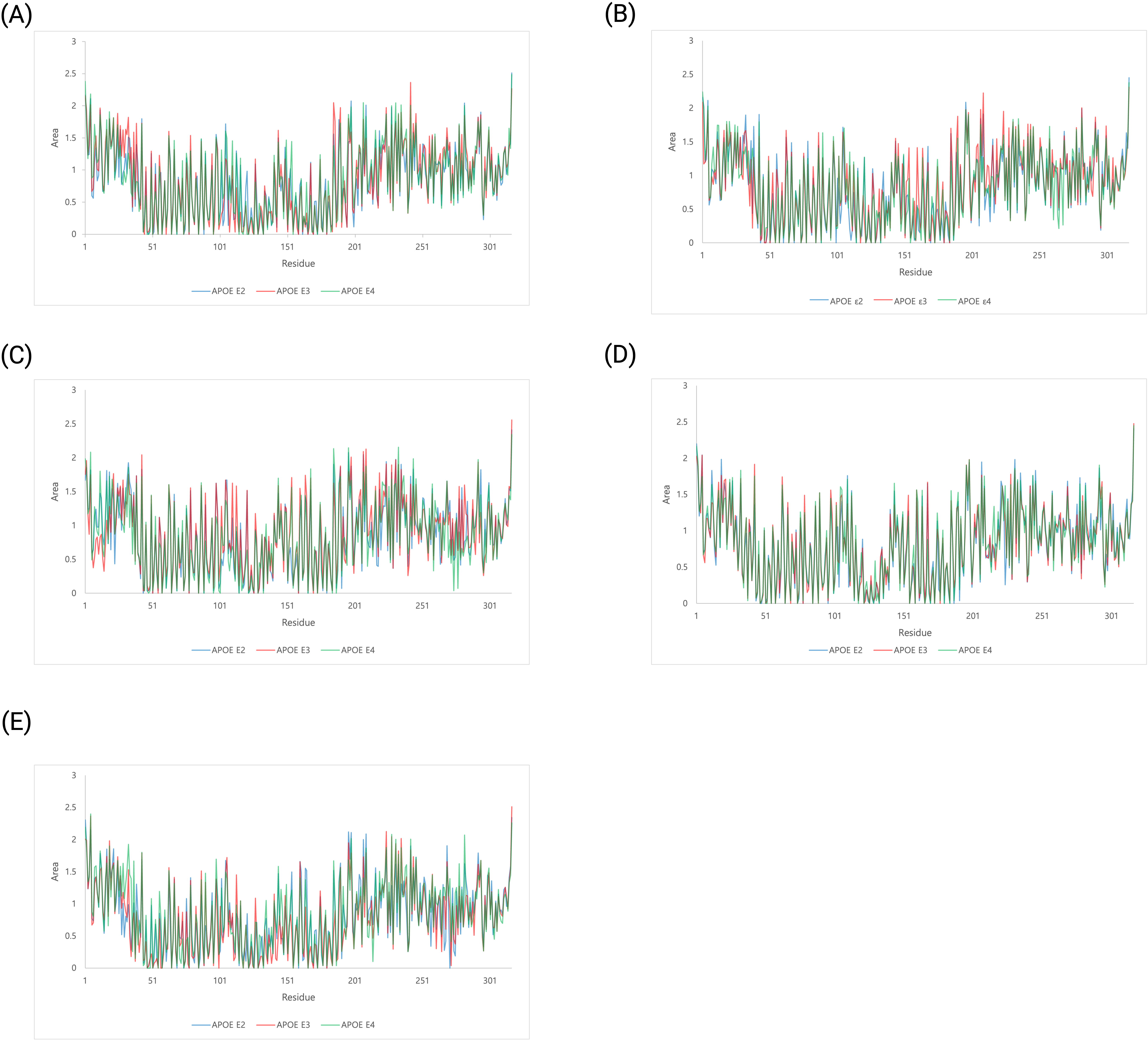

### Supplemental Fig.4.png

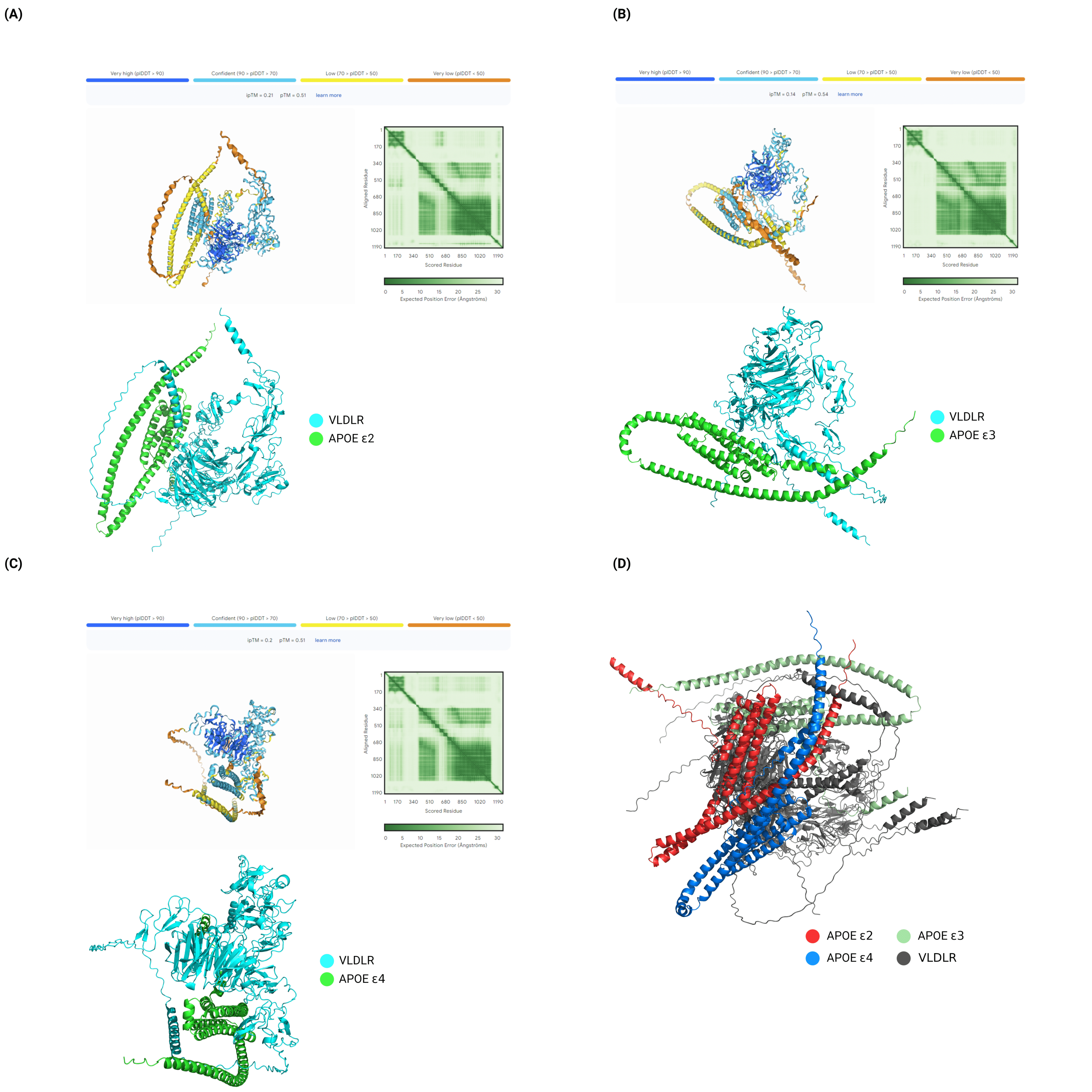

### Supplemental Fig.5.png

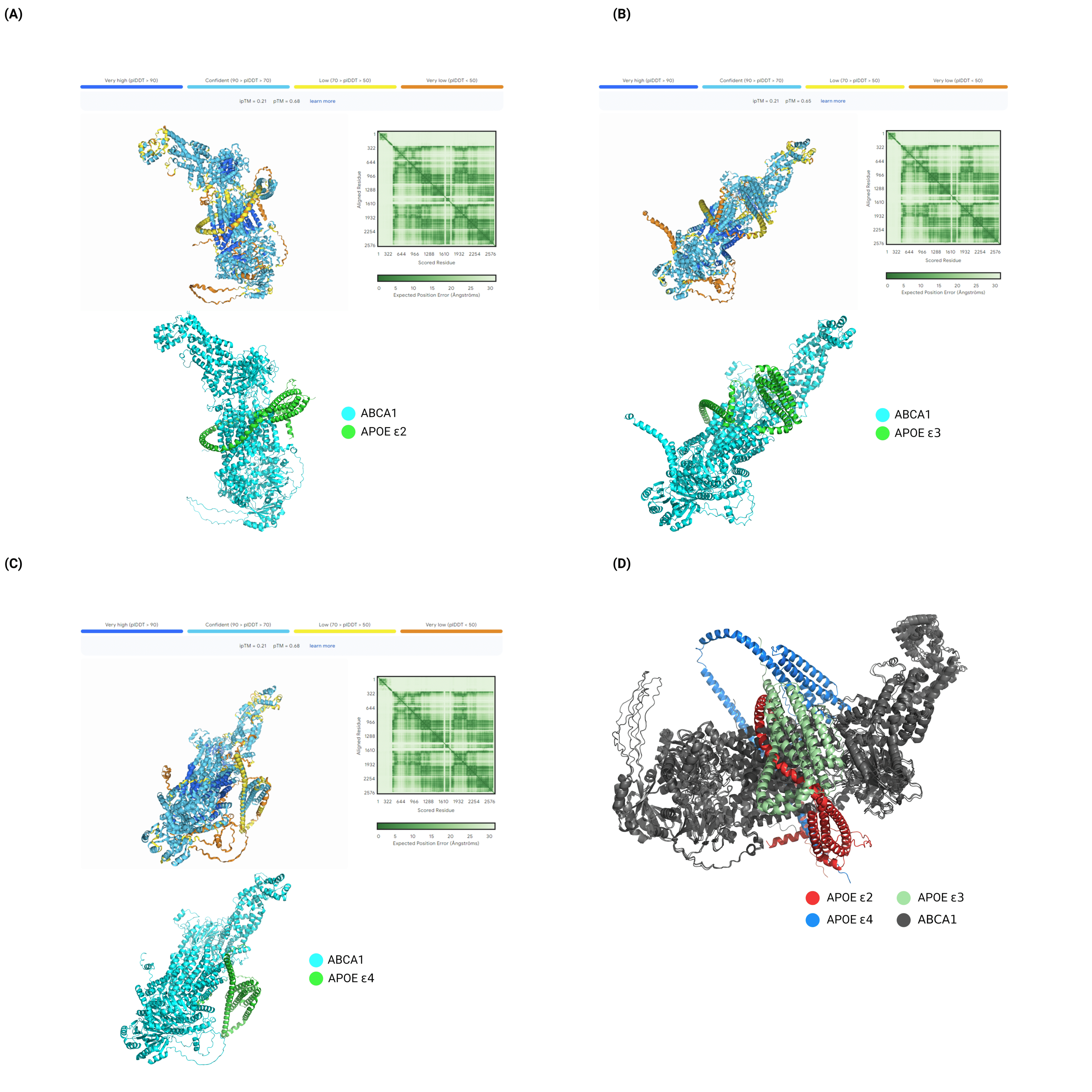

### Supplemental Fig.6.png

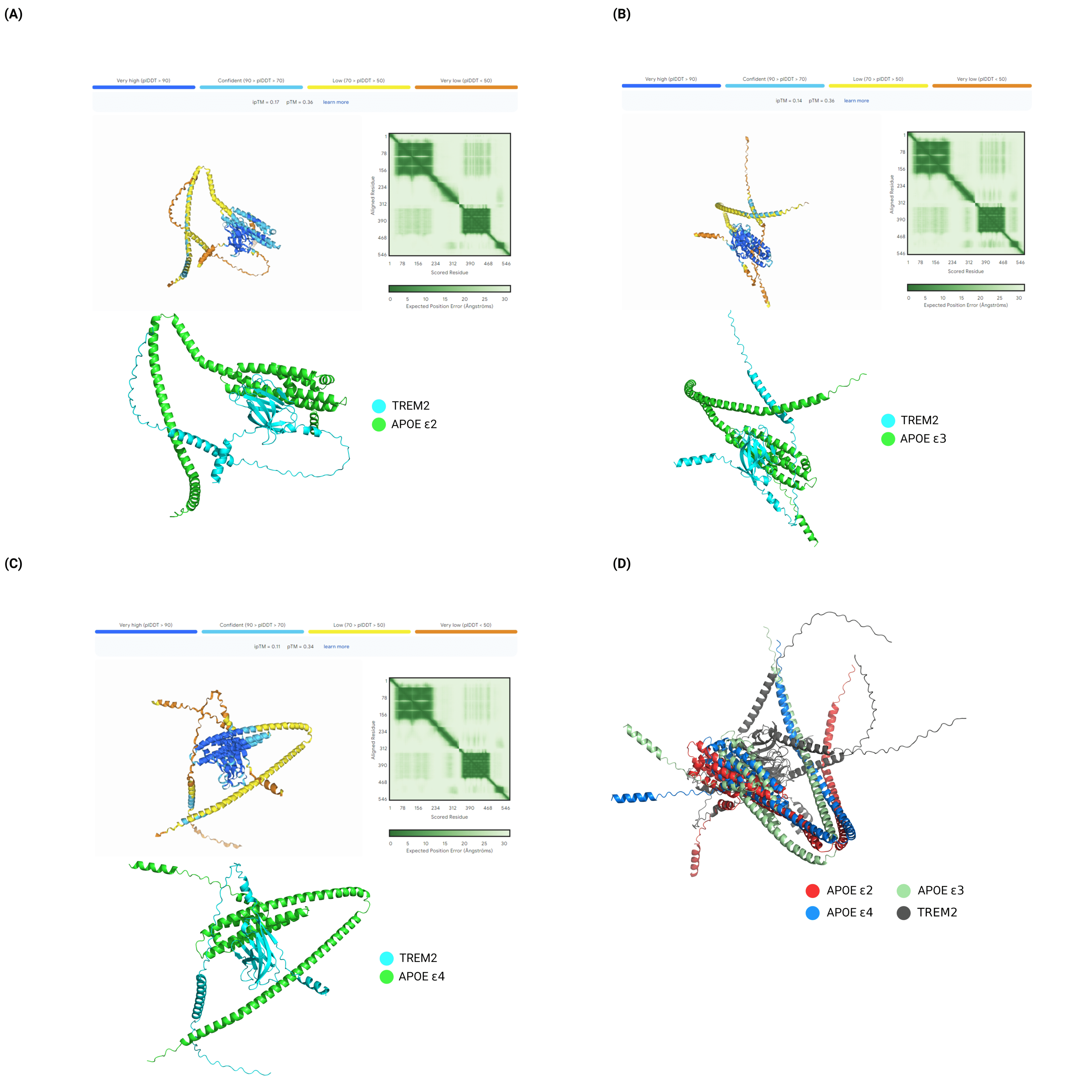
